# Characterising Structural Brain Connectivity of Patients with First Episode of Psychosis

**DOI:** 10.1101/2025.05.30.656096

**Authors:** Stella M. Sánchez, Thomas R. Knösche, Petra Fürstová, Antonín Škoch, Filip Španiel, Helmut Schmidt, Jaroslav Hlinka

## Abstract

**BACKGROUND AND HYPOTHESIS:** Schizophrenia is associated with widespread neuroanatomical abnormalities affecting both grey matter and white matter (WM). Early symptoms are often linked to dysfunctions in the frontal cortex and the temporal lobe. This study investigates WM disruptions and explores how structural connectivity (SC) may contribute to the underlying mechanisms of the disorder.

**STUDY DESIGN:** We analysed SC derived from diffusion MRI in 127 patients experiencing their first episode of schizophrenia (FES group), compared with healthy controls. Focusing on the fronto-parietal-temporal network, we examined SC across three hierarchical levels: network, node, and connection. SC metrics were compared between groups using 3-factor ANCOVA, accounting for relevant covariates. We also investigated associations between SC metrics and core positive symptoms using non-parametric correlations.

**RESULTS:** The FES group showed significantly reduced average SC strength and global efficiency within the fronto-parietal-temporal network. At the nodal level, SC strength was significantly lower in the left inferior and middle temporal gyri (L.ITG, L.MTG), and in the right inferior parietal gyrus (R.IPG) and temporal pole (R.TP). No significant group differences emerged at the connection level. Notably, SC strength in the R.IPG was negatively correlated with Conceptual Disorganisation scores.

**CONCLUSIONS:** Our findings reveal global and regional SC disruptions in early psychosis, particularly in areas supporting cognitive, language, and executive functions. The observed association between R.IPG connectivity and Conceptual Disorganisation supports the link between disrupted SC and formal thought disorder, reinforcing the role of impaired structural integration in early psychosis.

## 1. Introduction

**Schizophrenia** is a severe and chronic mental disorder characterised by recurrent psychotic episodes, affecting approximately 1 in 300 individuals worldwide^1^. The disorder typically manifests in late adolescence or early adulthood, with an earlier onset observed in males. Diagnosis is often established following the first psychotic episode, a critical window for early intervention.

The **first episode of psychosis (FEP)** is predominantly marked by positive symptoms, being the reason why the patient presents to the clinician. Among these, hallucinations—most commonly auditory but also visual, tactile, or olfactory—are the most frequently reported. Delusions, such as paranoia or grandiosity, and conceptual disorganisation, which manifests as disordered thinking and incoherent speech, are also core features. In addition to positive symptoms, negative symptoms—including diminished motivation, abulia, and reduced emotional expression—may also be present during the early stages. However, these symptoms occur less frequently than in chronic schizophrenia^2^ and are often underrecognised despite their significant impact on functional outcomes.

Identifying the **positive symptoms** mentioned above is crucial for accurate diagnosis and early intervention. According to the DSM-5, a diagnosis of schizophrenia requires the presence of at least two core symptoms—whether positive, negative, or cognitive—for a minimum duration of one month, with at least one symptom being hallucinations, delusions, or conceptual disorganisation^1^.

From a neuroanatomical perspective, schizophrenia is associated with widespread **structural alterations** affecting both grey matter (GM) and white matter (WM). The initial presentation of symptoms is strongly linked to dysfunctions in the frontal cortex and temporal lobe. Specifically, hallucinations are thought to originate in the temporal pole, while their persistence and severity are modulated by fronto-parietal-temporal network dysfunctions^3,4^.

**White matter alterations**, which reflect disrupted neuronal connectivity, can be investigated using **diffusion-weighted imaging** (DWI), an MRI-based technique sensitive to the Brownian motion of water molecules. Given that water diffusion in WM is constrained by fibre architecture, DWI enables the reconstruction of fibre trajectories and the quantification of **structural connectivity** (SC). Diverse studies, using metrics derived from DWI and SC, have reported WM alterations in schizophrenia patients^5^, as well as first-episode patients^6–8^, and early/late-onset schizophrenia and chronic patients^9^.

In this study, we conducted a comprehensive investigation of SC within the **fronto-parietal-temporal network** in a cohort of patients who had experienced their first episode of psychosis (FEP), a group often referred to as **first-episode schizophrenia (FES)**. To assess early-stage WM alterations, we performed a statistical comparison against a group of healthy controls (HC) with no history of psychosis. Our study aims to address the following key questions: 1) Do structural alterations in the fronto-parietal-temporal network emerge immediately after or during the FES, and can SC capture these changes?, 2) is it necessary to examine SC at different levels (from the coarse network level to the fine-grained connection level, considering also an intermediate node level) to fully evaluate its impact in this cohort?, and 3) are these connectivity alterations associated with early psychotic symptoms, such as auditory hallucinations? By addressing these questions, our study seeks to enhance the understanding of white matter abnormalities in early-stage schizophrenia and provide insights into the role of SC in its pathophysiology.

## 2. Methods

### 2.1. Participants

This study is based on the Early-Stage Schizophrenia Outcome (ESO) dataset, an ongoing large-scale prospective trial conducted in Prague and the Central Bohemia surveillance area. The ESO project investigates individuals experiencing a first episode of schizophrenia spectrum disorders and is coordinated by the National Institute of Mental Health (NIMH). The study was approved by the Ethics Committee of the Prague Psychiatric Center on June 29, 2011 (protocol code 69/11) and was conducted in accordance with the Declaration of Helsinki. All participants received a comprehensive explanation of the study procedures and provided written informed consent before enrollment. To ensure privacy and confidentiality, all sensitive participant data were anonymised.

For this study, we analysed data from the first visit (V1) of 127 individuals with first-episode of schizophrenia (FES). The diagnosis was confirmed by two experienced psychiatrists based on the ICD-10 Diagnostic Criteria for Research^10^. Inclusion criteria required that participants: (1) were undergoing their first psychiatric hospitalisation; (2) had an ICD-10 diagnosis of schizophrenia, acute and transient psychotic disorders, or schizoaffective disorders, as determined by the Mini-International Neuropsychiatric Interview (MINI^11^); (3) had experienced fewer than 24 months of untreated psychosis; and (4) were at least 18 years old. Individuals diagnosed with psychotic mood disorders, including schizoaffective disorder, bipolar disorder, or major depressive disorder with psychotic features, were excluded from the study.

A healthy control (HC) group was recruited through local advertisements, and 75 individuals were included. The main exclusion criteria for healthy controls were a personal history of any psychiatric disorder or substance abuse, as assessed using the MINI, and a family history of psychotic disorders in first- or second-degree relatives. Additional exclusion criteria for both FES and HC groups included current neurological disorders, a history of seizures or head injury with altered consciousness, intracranial haemorrhage or neurological sequelae, intellectual disability (IQ < 80), history of substance dependence, and any contraindication for MRI scanning^12^.

### 2.2. MRI Protocol and Clinical Data

MRI data were acquired with a 3T Prisma Siemens scanner.

<u>T1w</u> structural images were obtained by applying an MPRAGE sequence with the following parameters: TR/TE/TI = 2300/4.63/900 ms, flip angle = 10°, 162×210 voxels, voxel size = 1×1×1 mm^3^, FOV = 256 mm, 170 slices, matrix 256×256×224, bandwidth 130 Hz/pixel, GRAPPA acceleration factor 2 in phase-encoding direction, reference lines 32, prescan normalise on, elliptical filter on, raw filter off.

<u>DWI data</u>d were acquired by a spin-echo EPI sequence with TR/TE = 8300/84 ms, voxel size 2×2×2 mm^3^, flip angle = 90°, six b_0_ volumes in phase-encoding direction and six more in the reverse phase-encoding direction, b_1_ = 1000 s/mm^2^ and b_2_ = 2500 s/mm^2^ in 64 diffusion gradient directions each shell.

In addition, the participants underwent an intensive examination, including (but not limited to) neuropsychology testing and demographic assessments. The data used in this study are gender, age, years of education, and Positive and Negative Syndrome Scale (PANSS^13^) scores.

### 2.3. MRI Processing

#### 2.3.1. T1w processing

Although the primary objective of this study is to characterise and quantify differences in WM between groups, T1-weighted (T1w) anatomical images were essential to support specific steps of DWI processing and to complement the subsequent SC analysis.

Preprocessing of T1w images began with an initial visual inspection to detect and exclude images with major artefacts. Following this, several preprocessing steps were applied using tools from the FMRIB Software Library (FSL)^14,15^. These steps included: (1) coregistration of the T1w image to the DWI data using boundary-based registration (FLIRT-BBR); (2) skull stripping to remove non-brain areas using the Brain Extraction Tool (BET); and (3) tissue segmentation into different compartments using FMRIB’s Automated Segmentation Tool (FAST).

Subsequently, GM parcellation and T1w-derived metrics were obtained using the *recon-all* pipeline implemented in FreeSurfer. For further details on this procedure, refer to section 2.3 of Duarte-Abritta^16^.

#### 2.3.2. DWI processing

Preprocessing of DWI data began with a visual inspection to identify and exclude images with significant artefacts. Following this, a series of established correction steps was applied, including: (1) denoising to reduce signal fluctuations caused by thermal noise^17^; (2) removal of Gibbs ringing artifacts^18^; (3) correction of susceptibility-induced distortions; (4) eddy current and head motion correction; and (5) field inhomogeneity correction using the N4 algorithm implemented in ANTs^19^. The MRtrix3 software package^20^ was used for steps (1), (2), and (5), while TOPUP and EDDY from FSL were employed for susceptibility and motion correction^21,22^.

Following preprocessing, Fiber Orientation Distributions (FODs) were estimated using the multi-shell, multi-tissue (MSMT) Constrained Spherical Deconvolution (CSD) method^23^. To address residual intensity inhomogeneities, the intensity of WM and cerebrospinal fluid (CSF) components was normalised across all subjects^24^. The voxel-wise orientation information was then used to perform whole-brain probabilistic tractography using the Second-Order Integration over Fiber Orientation Distributions (iFOD2) algorithm, generating 10 million streamlines per subject^25^. The *seed-dynamic* option was employed to enhance streamline density distribution. To mitigate reconstruction biases and obtain a more biologically meaningful estimation of structural connection density, the streamlines were weighted using the Spherical Deconvolution Informed Filtering of Tractograms (SIFT2) algorithm^26^.

Further details on T1w and DWI processing can be found in the Supplementary Information of Sanchez et al.^27^.

### 2.4. Structural Connectivity Matrices

Two main elements are required to obtain an SC matrix per subject: a GM parcellation in each subject-native space and connectivity data between every pair of GM regions. For the first element, we used the Desikan-Killiany atlas^28^. This whole-brain atlas consists of 34 cortical areas, seven subcortical areas, and one cerebellum per hemisphere.

The second component, pairwise connectivity information, was derived from tractogram-based connectivity measures. Each tractogram consisted of streamlines generated using the probabilistic tractography algorithm (iFOD2). To enhance biological accuracy, streamline densities were subsequently reweighted to minimise discrepancies between the anatomical density of WM fibres and the density of reconstructed streamlines^26,29^.

Combining these two elements, we obtained an SC matrix per subject, where each entry (*C*_*ij*_) indicates the strength of the connectivity between regions *i* and *j*. Furthermore, each element *C*_*ij*_ is the sum of streamlines weighted^1^ (SSW)^26^. SC matrices are symmetric; in other words, the tractography algorithm is not capable of distinguishing the directionality of connections.

Based on the motivation from the Introduction, we focused on regions located in the frontal, temporal, and parietal lobes and their interconnections. According to the Desikan-Killiany atlas, each hemisphere contains 25 regions belonging to these lobes (see Supplementary Material for the complete list). We extracted these specific regions from the whole-brain SC matrices, yielding 50 × 50 SC matrices for further analysis.

To address the research questions outlined in the Introduction, we assessed SC matrices for the fronto-temporo-parietal network at three hierarchical levels: (1) the network level, analysing <u>average SC strength</u>, calculated as the average of all SC values in the matrix, and network efficiency, defined as <u>global efficiency</u> in graph theory^30,31^; (2) the node level, evaluating the <u>region of interest (ROI) strength</u>, defined as the sum of all SSW values associated with a given region; and (3) the connection level, comparing the <u>elements C_ij_</u>, where *i* or/and *j* is a region that arised significant at node level.

Additionally, we explored potential associations between SC variables—at any of the three levels described above—and the presence of auditory hallucinations, as assessed using item P3 of the PANSS scores. After finding no significant effect, we examined other positive core symptom items (P1: delusions and P2: conceptual disorganisation).

### 2.5. Statistical Analysis

To compare the distributions of demographic variables between the FES and HC groups, we first assessed normality using the Kolmogorov-Smirnov test. Depending on the distribution characteristics, group differences were then evaluated using either a t-test (for normally distributed variables) or a Mann-Whitney U-test (for non-normally distributed variables). For categorical variables, such as gender, we employed Fisher’s Exact Test to determine group differences.

Following prior literature (as elaborated in the Discussion section), we anticipated the necessity of controlling for potential confounding variables within the categories of demographic characteristics and T1w-derived metrics. Specifically, gender and intracranial volume (ICV) were identified a priori as confounding factors^32–34^. To address these, we employed analysis of covariance (ANCOVA), modelling SC as the dependent variable and adjusting for the identified predictors. When normal distributions might not be guaranteed, we applied nonparametric statistical tests for comparisons.

To examine associations between SC metrics and clinical variables, we performed Spearman’s correlation analysis, using a significance threshold of 0.05 to assess the strength and direction of relationships. Lastly, to account for multiple comparisons, we applied the Benjamini & Hochberg^35^ procedure to control the false discovery rate (FDR) and minimise the risk of Type I errors.

## 3. Results

### 3.1. Demographic and Clinical Data

Table 1 shows the demographic data of the sample and its comparison between groups. This study included 127 first-episode schizophrenia patients (FES) and 75 healthy control (HC) subjects. Although FES and HC were similar in age, statistical differences were found in gender (*p* = 0.0014) and years of education (*p* < 0.001). Table 1 reports (*mean* ± *SD)* (or frequency), as appropriate.

**Table 1.** Demographic and Clinical Characteristics of the Participants^2^.

|  | HC (n=75) | FES (n=127) | *p-value* | *Statistics* |
| --- | --- | --- | --- | --- |
| **Age (years)** | 30.22 ± 8.810 | 27.80 ± 7.014 | 0.0961 | 1.66 |

<sup>2</sup> Abbreviations: HC, Healthy Controls; FES, First-episode Schizophrenia; M/F, males/females; PANSS, Positive and Negative Syndrome Scale.

|  |  |  |  |  |
| --- | --- | --- | --- | --- |
| **Gender (M/F)** | 29/46 | 79/48 | 0.0014 | 0.3830<sup>3</sup> |
| **Education (years)** | 16.44 ± 2.78 | 14.66 ± 3.15 | < 0.001 | 4.057 |
| **PANSS** | - |  |  |  |
| **General** | - | 28 ± 7 |  |  |
| **Positive** | - | 11 ± 3.5 |  |  |
| **Negative** | - | 16 ± 5.8 |  |  |

All patients were drug-naive until the day of the MRI scanning, when they received antipsychotic medication.

### 3.2. ICV and Gender Analysis

To determine whether ICV might influence subsequent statistical analyses, we first assessed its distributional properties and whether it differed between groups.

Normality tests indicated that ICV values were compatible with a normal distribution in both groups (*p*_HC_ = 0.6287; *p*_FES_ = 0.6532). Based on these results, we performed an independent samples t-test to compare ICV between groups, which revealed no significant difference at the conventional alpha level of 0.05 (*p* = 0.0536). Given the proximity of this *p*-value to the threshold, we conducted a two-factor ANOVA to further investigate potential effects of group and gender on ICV.

The ANOVA results showed that ICV did not significantly differ between groups (*p*_group_ = 0.921), but was significantly different between males and females (*p*_gender_ << 0.001). No significant interaction was observed between group and gender (*p*_g:g_ = 0.198), indicating that the effect of gender on ICV was consistent across groups, and vice versa.

To further explore whether the gender effect on ICV was driving any group-level differences, we stratified the sample by gender and compared ICV distributions within each gender separately. No significant group differences were found within either males or females (*p*_M_ = 0.2338; *p*_F_ = 0.2187), suggesting that the overall gender effect observed in the ANOVA was likely due to an imbalance in gender composition between groups (HC and FES).

Based on these findings, and given the unequal gender distribution across groups, we included both gender and ICV as covariates in subsequent statistical analyses to control for their potential confounding effects.

### 3.3. Structural Connectivity Analysis

In this section, we present the outcomes of statistical analysis applied to the SC measures at the three explored levels. In all cases, we performed an ANCOVA with three predictors, group, gender and ICV, to elucidate whether any of these variables have an impact on the dependent variable (SC-derived metric) and, if so, which one.

For the <u>network level</u> analysis, we calculated two global metrics (average SC strength and network efficiency) and assessed them through a 3-factor ANCOVA (see Table 2). We found statistical differences in both metrics due to group, even after controlling for ICV. The top panel of Figure 1 displays the relation between the average SC strength and ICV, and the bottom panel shows the relation between Network Efficiency and ICV. In both cases, the relationship is positive and the HC group shows a higher mean than the FES group.

**Table 2.** SC Analysis at Network Level - Global Metrics^4^.

|  | GROUPS |  | 3-factor ANCOVA<sup>5</sup> |  |  |
| --- | --- | --- | --- | --- | --- |
| Metric | HC (n=75) | FES (n=127) | $p_{\text{group}}$ | $p_{\text{gender}}$ | $p_{\text{ICV}}$ |
| Average SC Strength | 17.98 ± 2.12 | 17.36 ± 2.56 | 0.0166 | 0.110 | 1.30e<sup>-8</sup> |
| Network Efficiency | 1.14 ± 0.142 | 1.09 ± 0.156 | 0.00227 | 0.0519 | 1.92e<sup>-10</sup> |

<sup>4</sup> Abbreviations: HC, Healthy Controls; FES, First-episode Schizophrenia; ANCOVA, Analysis of Covariance; SC, Structural Connectivity; ICV, Intracranial Volume.
<sup>5</sup> Outcomes from a 3-factor ANCOVA (setting group, gender, and ICV as factors). Reported p-values are uncorrected. Both metrics remain significant after FDR correction at $\alpha = 0.05$ .

**Figure 1.**
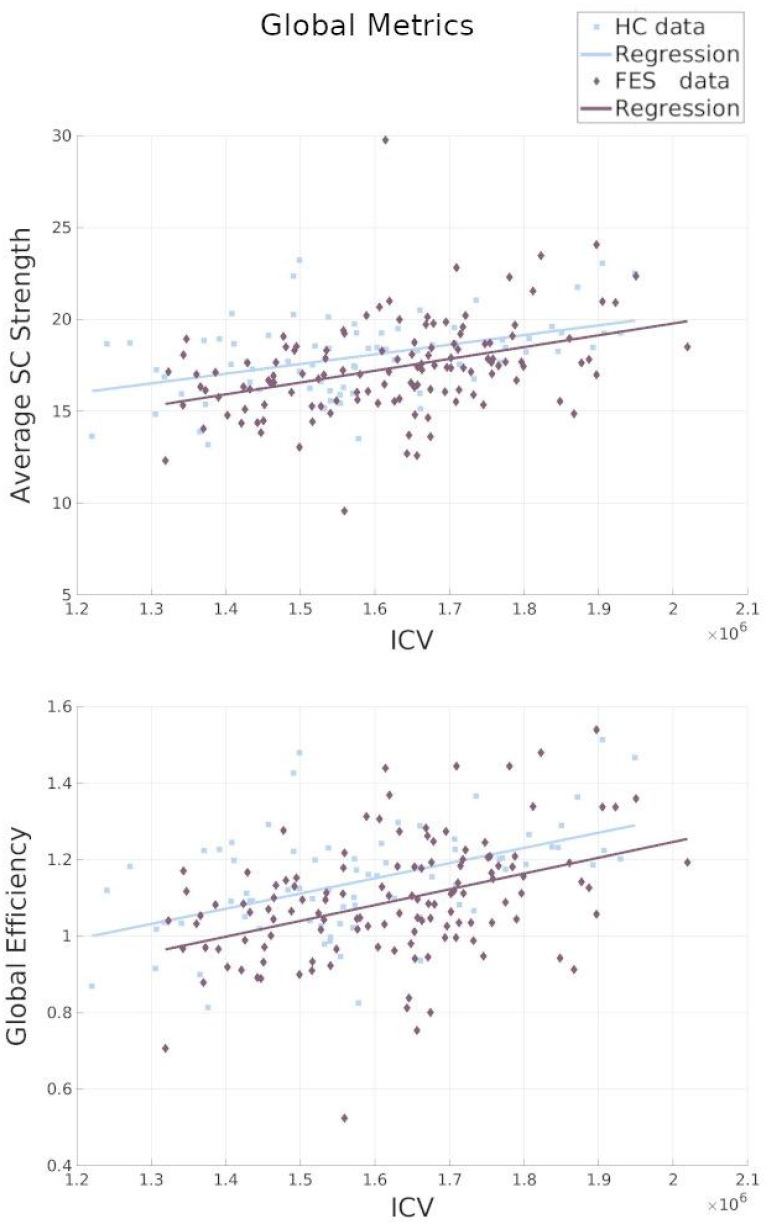
Global metrics plots. Scatter plots display HC (light blue squares) and FES (dark purple diamonds) data derived from SC. The solid lines represent the linear regression, only used for visualisation purposes. <u>Top pane</u>l: Average SC Strength vs ICV (mm^3^). From the regression model: HC group, β = 5.27 × 10^−7^, R^2^ = 0.20, p < 0.001; FES group, β = 6.43 × 10^−6^, R^2^ = 0.15, p < 0.001. <u>Bottom panel</u>: Global Efficiency vs ICV (mm^3^). From the regression model: HC group, β = 3.97 × 10^−7^, R^2^ = 0.25, p < 0.001; FES group, β = 4.13 × 10^−7^, R^2^ = 0.16, p < 0.001.

In addition, we examined whether ICV is related to the average SC strength and global efficiency. For all cases, we found significant p-values (<< 0.001) for both groups and positive (but moderated) Spearman’s coefficients: ρ_HC_ = 0.39 and ρ_FES_ = 0.42 for average SC strength; and ρ_HC_ = 0.47 and ρ_FES_ = 0.41 for global efficiency.

Regarding the <u>node level</u> (ROI strength analysis), we found significant differences (after FDR correction at α = 0.05) in the ROI Strength due to the *group effect* in four brain regions (displayed in Figure 2). This means that the group is a main predictor for these regions even after regressing out the factor ICV. In addition, these regions did not show statistical differences due to gender. Reported *p*-values are before applying FDR correction to *p*_group_ values (α = 0.05). Table 3 displays the outcomes from the 3-factor ANCOVA and shows that the HC group presents more ROI strength than the FES group.

**Table 3.** SC Analysis at Node Level - ROI Strength^6^.

| Hemisp. | ROI | GROUPS |  | 3-factor ANCOVA<sup>7</sup> |  |  |
| --- | --- | --- | --- | --- | --- | --- |
| | | HC (n=75) | FES (n=127) | $p_{group}$ | $p_{gender}$ | $p_{icv}$ |
| Left | ITG | 18.77 ± 3.52 | 17.46 ± 3.30 | 0.00115 | 0.948 | 0.000348 |
|  | MTG | 27.59 ± 3.85 | 26.24 ± 4.48 | 0.00380 | 0.851 | 2.09e<sup>-05</sup> |
| Right | IPG | 39.90 ± 5.82 | 37.68 ± 5.94 | 0.00245 | 0.809 | 0.00125 |
|  | TP | 3.611 ± 1.26 | 2.99 ± 1.18 | 0.000392 | 0.0272 | 0.00283 |

**Figure 2.**
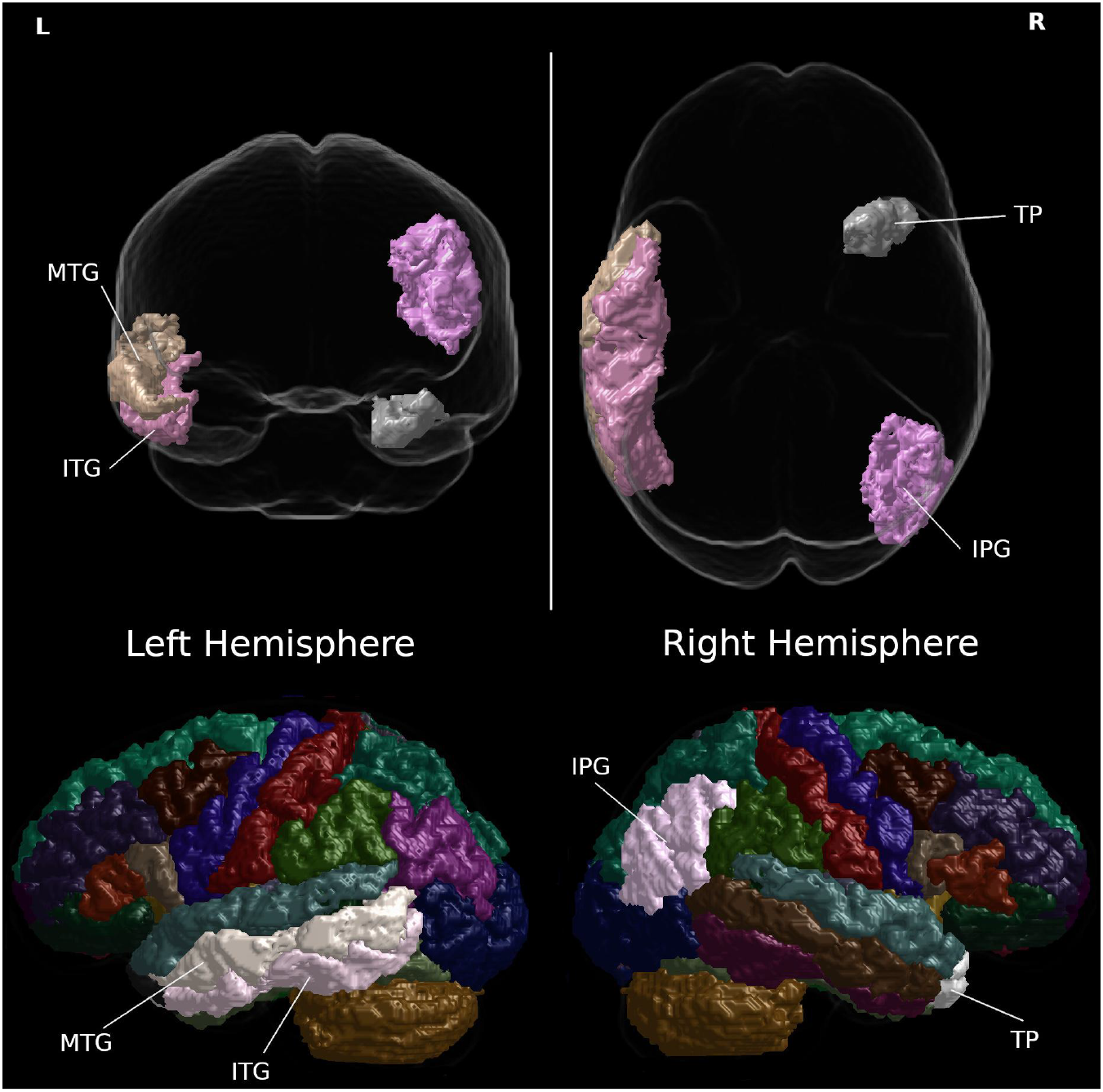
Grey Matter Regions with significant differences in SC Strength between groups. <u>Top panel</u>: Glass brain in coronal and axial views. L: left; R: right. <u>Bottom pane</u>l: regions highlighted in a whole-brain layout in sagittal view. The Desikan-Killiany (2006) atlas is used.

Figure 2 displays the brain regions from the Desikan-Killiany atlas that show significant differences between groups: the Inferior Temporal Gyrus (ITG) and the Middle Temporal Gyrus (MTG) from the left hemisphere, and the Inferior Parietal Gyrus (IPG) and the Temporal Pole (TP) from the right hemisphere.

Figure 3 shows four panels (one for each ROI listed in Table 3). Each panel illustrates the relationship between ICV and the ROI strength (dependent variable). The scatter plots display the observed data points for each group, while the solid lines are regression lines. Only the right TP shows a negative relationship between the strength of the ROI and ICV in the FES group, but not in HC. In all cases, the estimated mean for the HC group was higher than that of the FES group.

**Figure 3.**
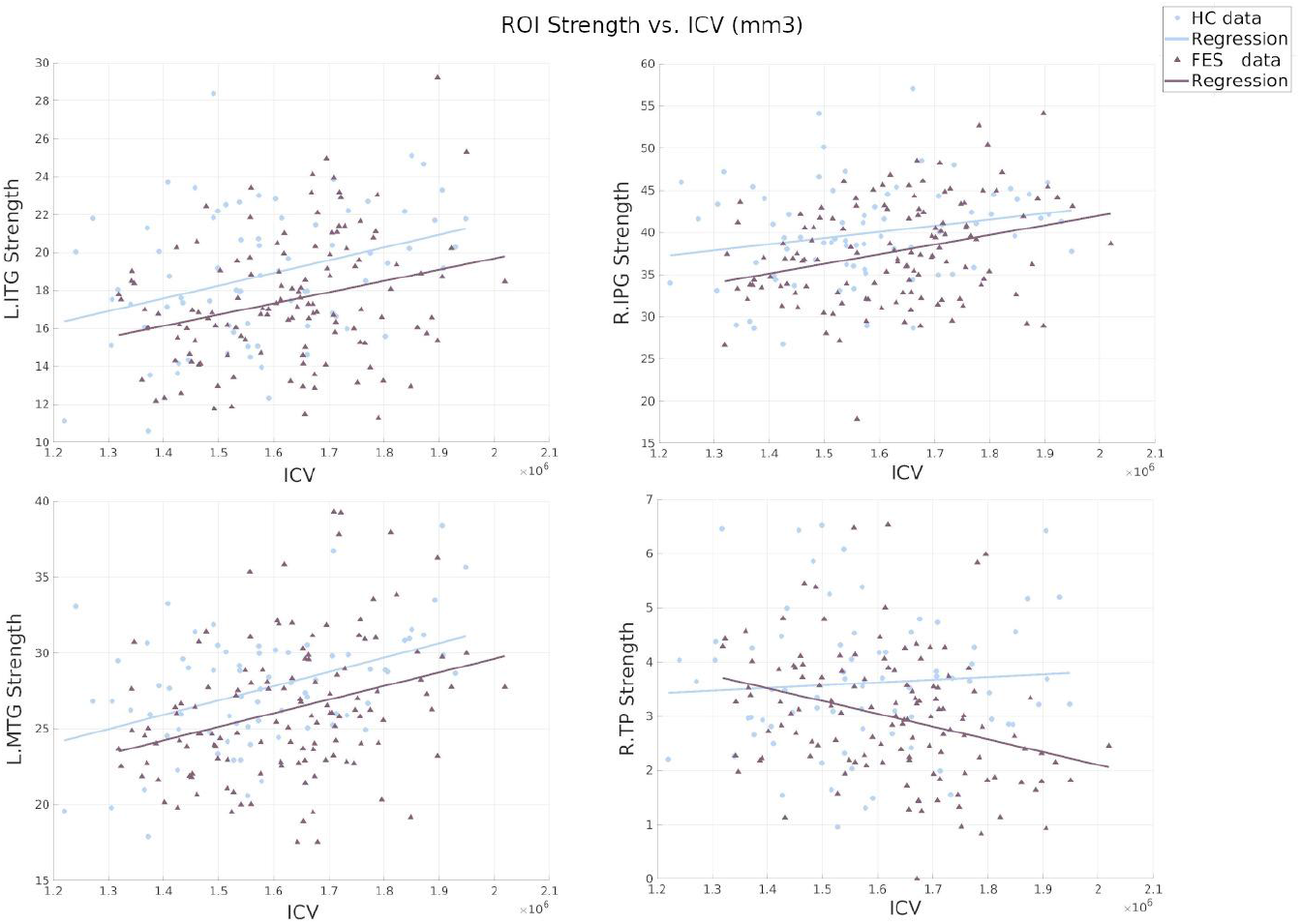
Relationship between the ROI strength and ICV (mm^3^). Scatter plots display the HC (light blue circles) and FES (dark purple triangles) data derived from SC. The solid lines are regression lines.

No statistical differences were found at the <u>connection level</u> analysis. As mentioned in the Discussion section, there are two main reasons for this: (1) the FES group may not differ structurally from the control group at a fine-grained level, and (2) the statistical power may be limited due to the large number of hypotheses tested at this level of analysis.

### 3.4. Structural Connectivity and Clinical Data

We sought an association between the specific subcategory of the PANSS corresponding to hallucinations and the SC strength of ROIs listed in Table 3. Unexpectedly, we found no significant association that survives FDR correction when applying Spearman’s correlation (with right IPG: *p* = 0.039, ρ = -0.189). However, we found a negative and statistical correlation between the SC-strength of the right IPG and the second positive (P2) PANSS subcategory corresponding to ‘Conceptual Disorganization’ (ρ = -0.288, *p* = 0.00100, p_FDR_ = 0.022). Figure 5 displays the corresponding scatter plot for this last finding.

**Figure 5.**
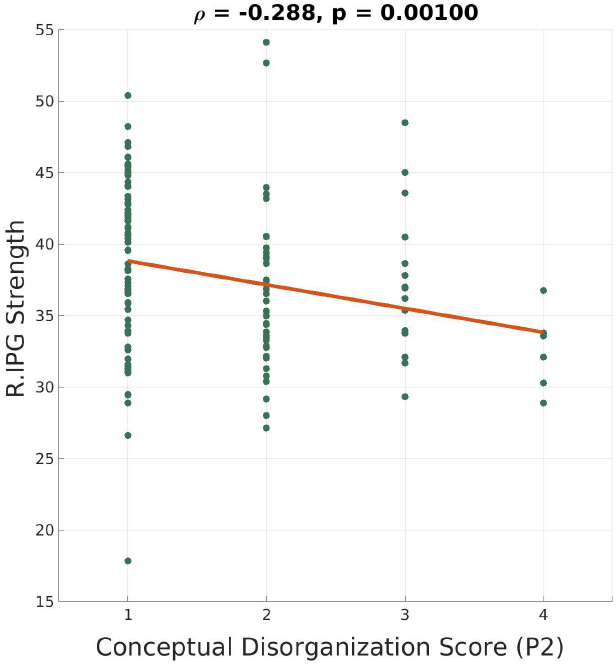
Scatter plot displaying the relationship between the SC Strength of R.IPG and the scores for the subcategory ‘Conceptual Disorganization’ (P2) of PANSS. Dark green dots are the data for FES group. A regression line (orange) is included to visualise the trend. The negative correlation (ρ=−0.288) suggests a weak inverse association, with a statistically significant p-value (p = 0.00100).

## 4. Discussion

In this study, we aimed to investigate whether a first episode of schizophrenia (FES) leaves detectable alterations in white matter (WM) and whether structural connectivity (SC) can reveal these changes. To achieve a comprehensive evaluation of SC derived from diffusion-weighted imaging (DWI), we analysed three hierarchical levels: network, node and connection. Each level provides a distinct perspective, from coarse-grained global patterns to fine-grained local changes. Based on existing literature^1,3^, we focused on brain regions implicated in the early manifestation of schizophrenia. Additionally, we examined potential associations between SC metrics and Positive and Negative Syndrome Scale (PANSS) scores, particularly those related to the first symptoms of schizophrenia.

Before addressing the research questions outlined in the Introduction, we first examined the structure of our data sample. We found that the first-episode schizophrenia (FES) group was comparable in age to the healthy control (HC) group. As expected based on previous literature^36,37^, the FES group exhibited significantly fewer years of education than the HC group, which may be related to early cognitive and functional difficulties associated with the disorder. Additionally, we observed a gender imbalance in our sample, consistent with previous findings that schizophrenia samples tend to include a higher proportion of male patients^32^. In such cases, two methodological approaches are typically considered: (1) matching the control group by gender, or (2) including gender as a covariate in statistical analyses. Moreover, the literature has long debated the potential influence of gender on the onset and manifestation of schizophrenia symptoms, whether positive or negative^38,39^.

Beyond these demographic considerations, we also examined the potential systematic effects related to brain size, particularly relevant for tractography and the construction of structural connectomes. It has been recommended to account for intracranial volume (ICV) as a nuisance variable to minimise bias in the reconstruction and comparison of WM fibres^34^. Controlling for ICV also facilitates clearer and more interpretable results^34^.

Taking these factors into account, and to preserve statistical power, we chose not to reduce the sample size in order to achieve gender balance. Instead, we applied a robust statistical framework to examine the relationships between subject group, gender, and ICV. Our analyses indicated that ICV is independent of group status but is significantly influenced by gender. As the observed ICV differences appeared to be driven by the gender imbalance, and we could not fully disentangle the effects of gender and ICV, both variables were included as covariates in subsequent statistical analyses to account for their potential confounding effects^32–34^.

To address questions 1 and 2, we evaluated SC at the global network level by assessing two graph-theoretical measures: average SC strength and global efficiency within the fronto-parietal-temporal network. Both metrics were significantly lower in FES compared to HC, even after regressing out gender and ICV effects. These results suggest that patients with FES exhibit reduced network integration and disrupted global topology, which aligns with previous findings in schizophrenia^40^ and drug-naïve patients^41^. As proposed in earlier work^42^, these reductions in global SC metrics may be attributed to decreased interregional connectivity or disconnection along long-range pathways.

The following step in our investigation was to understand which nodes or connections contribute to these differences at network level. At the node level, we found significant differences in the ROI strength metric in four brain regions, even after accounting for gender and ICV as covariates, confirming that these differences can be attributed to the group effect. Within the fronto-parietal-temporal network, the following brain regions exhibited significantly lower SC strength in FES patients compared to HC: inferior and middle temporal gyri (ITG and MTG) from the left hemisphere, inferior parietal gyrus (IPG) and temporal pole (TP) from the right hemisphere. These regions are functionally associated with key cognitive, social, and sensory processes, many of which are affected in schizophrenia. The TP is heavily involved in social and emotional processing, as well as multisensory integration (auditory, visual, and olfactory)^43^, and has frequently been reported as structurally and functionally altered in schizophrenia^44^. The MTG plays a role in language processing and semantic memory, while the ITG is crucial for visual perception and sensory integration—both functions known to be disrupted in schizophrenia^45^. The right IPG is associated with attentional processing, social cognition, and body perception^46^.

Finally, we examined the finest scale of our SC metrics to identify specific connections that might contribute to the observed ROI strength differences between groups. We focused on connections involving at least one region identified in the node level analysis. However, no significant group differences emerged at the connection level. This likely reflects a lack of statistical power, given the high specificity of this analysis and the large number of comparisons involved.

We hypothesised an association between SC metrics and the PANSS hallucinations score (P3). However, no significant relationships were observed, contradicting our initial expectations (question 3). Given that at least one of the core positive symptoms must be present for a schizophrenia diagnosis (DSM-5), we expanded our analysis to PANSS P1 (delusions) and P2 (conceptual disorganisation). Interestingly, we identified a significant negative correlation between SC strength in the right IPG and P2 scores, indicating that lower SC strength in this region is associated with greater conceptual disorganisation.

This finding aligns with longstanding evidence linking formal thought and speech disorders to schizophrenia’s disorganisation dimension^47^. Conceptual disorganisation encompasses symptoms such as tangentiality, derailment, incoherence, poverty of content of speech, and difficulty in abstract thinking—all of which are hallmarks of schizophrenia’s cognitive dysfunction^47^. Furthermore, the right IPG has been implicated in conceptual processing and abstract thinking, particularly in numerical cognition^48^. The observed association suggests that disruptions in right IPG connectivity may underlie deficits in abstract thinking and speech organisation in schizophrenia patients, warranting further investigation in future studies.

In summary, our findings suggest that while connection level SC does not show alterations in FES, most likely due to a lack of statistical power, node and network level disruptions are evident, particularly in regions supporting cognitive, language, and executive functions. The observed decrease in network integration and the association between right IPG SC and conceptual disorganisation provide further evidence for disrupted structural connectivity in early schizophrenia. These findings highlight the need for further studies to replicate these results and explore their implications for early intervention and treatment strategies in psychosis.

## Acknowledgments

The authors would like to sincerely thank all subjects for their time and effort.

## Ethical statement

The authors confirm that all experiments involving human participants were performed following relevant guidelines and regulations. The study was conducted according to the guidelines of the Declaration of Helsinki and approved by the Ethics Committee of the Prague Psychiatric Centre (protocol code 69/11, approved on 29 June 2011). Written informed consent was obtained from all subjects involved in the study.

## Data Availability Statement

Freesurfer, FSL and MRtrix3 open-sourced softwares were implemented to process and analyse T1 and DWI data. The dataset analysed during the current study is not publicly available as it contains patients’ personal information that cannot be publicly shared due to the hospital policy, but is available from the corresponding author on reasonable request.

## Footnotes

1 The weights of each streamline connecting a pair of regions are summed to give the weight of that edge.

2 Abbreviations: HC, Healthy Controls; FES, First-episode Schizophrenia; M/F, males/females; PANSS, Positive and Negative Syndrome Scale.

3 Fisher’s Exact Test.

4 Abbreviations: HC, Healthy Controls; FES, First-episode Schizophrenia; ANCOVA, Analysis of Covariance; SC, Structural Connectivity; ICV, Intracranial Volume.

5 Outcomes from a 3-factor ANCOVA (setting group, gender, and ICV as factors). Reported p-values are uncorrected. Both metrics remain significant after FDR correction at α = 0.05.

6 Abbreviations: HC, Healthy Controls; FES, First-episode Schizophrenia; ANCOVA, Analysis of Covariance; ICV, Intracranial Volume; ROI, Region of Interest; ITG, Inferior Temporal Gyrus; MTG, Middle Temporal Gyrus; IPG, Inferior Parietal Gyrus; TP, Temporal Pole.

7 Outcome from a 3-factor ANCOVA (setting group, gender, and ICV as factors). P-values reported are uncorrected. The four brain regions present significant differences due to group after FDR correction at α = 0.05.

