## Supplemental Material for "Characterising Structural Brain Connectivity of Patients with First Episode of Psychosis"

### Supplementary Material

**Methods**

Brain Regions selected from Desikan-Killiany Atlas (2006) to build the fronto-parietal-temporal network:

Frontal Pole

- Caudal Middle Frontal Gyrus (CMFG)
- Lateral Orbito Frontal Gyrus (LOFG)
- Medial Orbito Frontal Gyrus (MOFG)
- Paracentral Gyrus (PaCG)
- Pars Opercularis (POP)
- Pars Orbitalis (POR)
- Pars Triangularis (PTR)
- Precentral Gyrus (PrCG)
- Rostral Middle Frontal Gyrus (RMFG)
- Superior Frontal Gyrus (SFG)
- Frontal Pole (FP)

Parietal Pole

- Inferior Parietal Gyrus (IFG)
- Postcentral Gyrus (PoCG)
- Precuneus (PCU)
- Superior Parietal Gyrus (SPG)
- Supramarginal Gyrus (SMG)

Temporal Pole

- Banks STS (BSTS)
- Entorhinal Cortex (EC)
- Fusiform Gyrus (FG)
- Inferior Temporal Gyrus (ITG)
- Middle Temporal Gyrus (MTG)
- Parahippocampal Gyrus (PHIG)
- Superior Temporal Gyrus (STG)
- Temporal Pole (TP)
- Transverse Temporal Gyrus (TTG)

**Results**

Coefficients from regression models for Figure 3:

| | $\beta$ | $R^2$ | $p$ |
|---|---|---|---|
| L.ITG | | | |
| HC | $6.72 \times 10^{-6}$ | 0.12 | 0.0029 |
| FES | $5.93 \times 10^{-6}$ | 0.076 | 0.0017 |
| L.MTG | | | |
| HC | $9.42 \times 10^{-6}$ | 0.19 | < 0.001 |
| FES | $8.96 \times 10^{-6}$ | 0.094 | < 0.001 |
| R.IPG | | | |
| HC | $7.33 \times 10^{-6}$ | 0.050 | 0.05 |
| FES | $1.15 \times 10^{-5}$ | 0.087 | < 0.001 |
| R.TP | | | |
| HC | $5.01 \times 10^{-7}$ | 0.005 | 0.55 |
| FES | $-2.35 \times 10^{-6}$ | 0.094 | < 0.001 |

**Supp. Table 1:** *Coefficients from the regression model illustrated in Figure 3.*
